# Metagenomic insights for antimicrobial resistance surveillance in soils with different land uses in Brazil

**DOI:** 10.1101/2022.12.05.519117

**Authors:** João Vitor Wagner Ordine, Gabrielle Messias de Souza, Gustavo Tamasco, Stela Virgilio, Ana Flávia Tonelli Fernandes, Rafael Silva-Rocha, María-Eugenia Guazzaroni

## Abstract

Anthropization in terrestrial environments commonly leads to land use transformation, changing soil properties and their microbial communities. This, combined with the exacerbated use of antibiotics in human and animal health promotes the expansion of the soil resistome. Considering the urgent need for surveillance of antimicrobial resistance (AMR), we aimed to evaluate how different land practices (urban, farming and forest) can affect the soil resistome and the dynamics of their bacterial communities. We collected eight soil samples from different locations in the countryside of São Paulo (Brazil), assessed the community profiles based on 16S rRNA sequencing and analyzed the soil metagenomes based on shotgun sequencing. Our results highlight differences in the communities’ structure and their dynamics which were correlated with land practices. Additionally, differences were observed in the abundance and diversity of antibiotic resistance genes (ARGs) and virulence factors (VFs) across studied soils, where a higher presence and homogeneity of *vanRO, mtrA* and *rbpA* genes were detected in livestock soils. We observed that *Staphylococcus* and *Bacillus* are positively correlated with each other and are markers for agricultural communities. Moreover, the abundance and diversity of ARGs and VFs observed in farming soils raises concerns regarding the potential spread of these genes in the environment. Together, our findings reinforce the importance and urgency of AMR surveillance in the environment, especially in soils undergoing deep land use transformations due to anthropic activity.

## 1. Introduction

Antimicrobial resistance (AMR), one of the most serious health risks of the 21st century, is a common competition mechanism used by environmental bacteria to ensure their survival in their natural environment. Although evidence of antimicrobial resistance in bacteria dates back to the pre-antibiotic era, studies suggest that human activity has a significant impact on the extension and diversity of the bacterial resistome [1–3].

Even though the soil microbiota naturally presents a large and robust diversity of ARGs in its intrinsic resistome, land use transformation due to anthropic activities, such as the excessive use of antibiotics in livestock production [4], antibiotic-enriched manure application [5,6], and excessive use of xenobiotics in crops [7,8], along with increasing levels of deforestation for farming or urban purposes, can alter bacterial communities and disseminate ARGs throughout the environment [9]. In these highly modified sites, soil bacteria more frequently are in close proximity to commensal and pathogenic ones, which could lead to an increased horizontal ARG transfer rate among them, followed by a dominance of organisms with acquired resistance in comparison to intrinsically resistant bacteria [10].

The “One Health” surveillance approach takes into consideration the interrelated link among people, non-human animals, and the environment [2]. The substantial role of the latter in AMR spread can be noted in the soil’s capacity to serve as a resistance gene reservoir, facilitating the spread of ARGs found in mobile genetic elements (MGEs), such as plasmids, integrons, and transposons, among different bacterial species, speeding the development of multidrug-resistant (MDR) pathogens [11]. According to Gosh et al. (2021) [12], MDR pathogens are thought to be responsible for up to 10 million cases of fatalities annually worldwide, with a mortality rate of 392,000 in Latin America.

Brazil, the largest country in Latin America, presents a population of 192 million people. Although the National Health Regulatory Agency (ANVISA) in Brazil has compiled data regarding healthcare-associated infections and levels of antimicrobial resistance on clinical settings over the last decades [13], the country is not equipped with a central microbiology reference laboratory, increasing the difficulty to conduct national data analysis regarding bacterial resistance [14]. The São Paulo state, located in the southeastern region in Brazil, inhabited by over 45 million people [15], is a critical region in terms of high levels of resistance among important pathogens such as non-fermenting Gram-negative bacilli and Gram-positive cocci, as *Staphylococcus aureus*. [15].

Recent estimates suggest that Brazil was responsible for almost 8% of all antibiotic consumption for veterinary purposes globally in 2017 and has an increased consumption projection of 11.8% in 2030 [16]. This is mainly due to a shift toward intensified livestock production systems that regularly use antimicrobial agents, which can directly affect the number of ARGs disseminated through the environment and, consequently, might contribute to increased levels of AMR in clinical settings [16].

Previous studies have shown the seriousness and urgent need to tackle AMR in the countryside of São Paulo, Brazil, especially after COVID-19 pandemic [17]. During this period, high rates of antibiotic use in hospitalized patients and prolonged time in invasive therapy have caused an alarming increase in polymyxin B-resistant *Klebsiella pneumoniae* isolates in 2021 [18] and a nosocomial outbreak of extensively drug-resistant (EDR) *K. pneumoniae* in 2022 [19].

Despite the extreme importance of monitoring healthcare infections associated with antibiotic resistant bacteria to combat AMR, there’s still a lack of data regarding environmental levels of resistance in northeastern soils of the São Paulo state, where there has been a historical land use of soils for urban construction, agricultural, and livestock practices, leading to a population density of approximately 1.75 million inhabitants [20–22]. Considering the urgent need to tackle AMR not only on clinical settings, but also taking into account the One Health approach, we used 16S rDNA and shotgun sequencing to profile the bacterial communities and resistomes of eight sites in the countryside of São Paulo, to provide the first, to our knowledge, environmental AMR surveillance of soil samples in this region.

## 2. Materials and Methods

### 2.1. Study area and sample collection

The samples used in this study were collected from different sites across São Paulo’s northeast region (Figure S1 and Table 1). Approximately 50 g of samples were aseptically taken from the upper 10 cm layer, after a 5 cm removal of litterfall, and placed in sterile Falcon tubes. For each selected site, 3 samples were collected and mixed for a better representation of the microbial community within. In total, eight samples were collected from: (i) Permanent Preservation Area (PPA) in the University of São Paulo campus - Ribeirão Preto, São Paulo; (ii) University of São Paulo campus lawn - Ribeirão Preto, São Paulo; (iii) Agriculture site - Sertãozinho, São Paulo; (iv) Pasture field - São Carlos, São Paulo; (v) Livestock site - Taquaritinga, São Paulo; (vi) Hen house - São Carlos, São Paulo; (vii) Orchard field - São Carlos, São Paulo and (viii) Urban square - Sertãozinho, São Paulo.

**Table 1.** Main characteristics and soil sample’s locations.

| Soil Samples | Geological Classification | Land Use Classification | Anthropic Activity | Sampling Site | Coordinates |
| --- | --- | --- | --- | --- | --- |
| Agriculture (Agri) | Red Latosol | Farming | Large sugarcane crop site + Agrochemical use | Sertãozinho, SP | -21.16643, -47.99004 |
| Pasture (Pas) | Deep Quartz Sand | Farming | Small familiar cattle farm | São Carlos, SP | -21.921, -47.90373 |
| Cattle Site (Catt) | Bauru Sandstone | Farming | Large cattle site. High circulation of people and livestock | Taquaritinga, SP | -21.48066, -48.54118 |
| Orchard (Orch) | Deep Quartz Sand | Farming | Small familiar orchard site | São Carlos, SP | -21.92674, -47.87008 |
| Hen House (Hen) | Deep Quartz Sand | Farming | Small familiar poultry farm | São Carlos, SP | -21.95133, -47.89079 |
| Urban Square (UrbSq) | Red Latosol | Urban | High circulation of people and small animals | Sertãozinho, SP | -21.1134, -47.98762 |
| USP* Permanent Protection Area (PPA) | Red Latosol | Forest | Protected area. Access restricted to research | Ribeirão Preto, SP (USP) | -21.1662, -47.86036 |
| USP Campus Lawn (Lawn) | Red Latosol | Forest | High circulation of people and animals | Ribeirão Preto, SP (USP) | -21.16511, -47.85944 |

\* University of São Paulo (USP)

All soils were allocated in three categories, according to their land uses over the last decades, being those (i) Farming - sugarcane field, livestock site, hen house, pasture field and orchard; (ii) Urban - urban square and (iii) Forest - PPA and campus lawn.

### 2.2. DNA extraction and sequencing

The metagenomic DNA of each soil was extracted using DNAeasy Powersoil^®^ Kit (QIAGEN), following manufacturer’s recommendations. The quantification and quality analysis of the extracted DNA was performed using Nanodrop™ One (Thermo Fisher Scientific) and by an agarose (1%) electrophoresis gel. Part of the extracted DNA was used for triplicates amplification of 16S rDNA, following Nanopore - 16S Barcoding Kit 1-24 (SQK-16S024, Oxford Nanopore Technologies - ONT) recommendations, adding different barcodes for each replicate. All samples were individually purified, quantified by Nanodrop™ One and mixed in proportioned amounts in order to make a representative pool of all soil samples. A single multiplex sequencing was performed using the aforementioned kit and Flongle flowcells in MinION model Mk1B. The remaining extracted DNA was submitted to metagenomic sequencing on Illumina NovaSeq 6000 platform at Novogene (Sacramento, CA, USA), with sequencing depth of 12 Gb/sample. Table S1 shows the data quality summary of raw data from shotgun sequencing.

### 2.3. Data processing and analysis

#### 2.3.1. Amplicon sequencing

16S rDNA reads’ processing and analysis were performed as de Siqueira and collaborators (2021) [23]. Briefly, reads were base-called using Guppy Basecalling Software (version 6.1.3) with dna_r9.4.1_450bps_hac.cfg configuration file [24]. Base-called reads had their quality assessed by NanoStat (version 1.6) and NanoFilt (version 2.8) was used to select reads with quality scores above Q7 [25]. After the initial filtering step, demultiplexing of reads was performed by Porechop (version 2.4) using the barcodes from 16S Barcoding Kit 1-24 (SQK-16S024). Demultiplexed reads were mapped to a 16S rDNA NCBI reference database using minimap2 (version 2.17) [26]. All further analyses and data plotting were performed in R (version 4.1.0), using tools in the package Vegan (version 2.5.7) [27].

#### 2.3.2. Shotgun sequencing

Shotgun sequencing raw data was processed with the fastp (version 23.1) tool (https://github.com/OpenGene/fastp), for adapter and low-quality reads removal [28]. High quality reads assembly was carried out with the MegaHIT tool (https://github.com/voutcn/megahit) and the metagenome annotation was performed with Prokka tool (https://github.com/tseemann/prokka) [29], with the assembly statistics calculated with the assembly_stats tool (https://github.com/sanger-pathogens/assembly-stats). The identification of ARGs, virulence factors (VFs) and plasmid markers was performed with the ABRICATE pipeline (https://github.com/tseemann/abricate), with an identity cut-off of 80%, by searching previously annotated genes in reference databases (ARG-ANNOT, CARD, PlasmidFinder, ResFinder and VFDB) [30–34].

## 3. Results

### 3.1. Bacterial Community Dynamics

To understand the differences in bacterial communities of soils from different locations and land uses, we collected eight soils from different sites across São Paulo’s northeast region (Table 1 and Figure S1), and we assessed the community profiles based on 16S rRNA sequencing trough Oxford Nanopore MinION device, which allows full-length sequencing of the 16S gene (Figure 1A). In all sampling sites, *Bacillus* was observed as one of the most ubiquitous bacteria in the communities, with relative abundances of 1.7% in campus lawn soils and in PPA, 2% in urban square, 2.4% in orchard, 4.4% in hen house, 11.7% in cattle sites, 17.7% in pasture area and 31.1% in agriculture soils. Two other frequent bacteria genera identified were *Vicinamibacter*, with relative abundance ranging from 1.9% (orchard) to 6.2% (pasture) and *Rhodoplanes*, ranging from 1.9% (hen house) to 4.4% (PPA soil) (Table S2).

**Figure 1.**
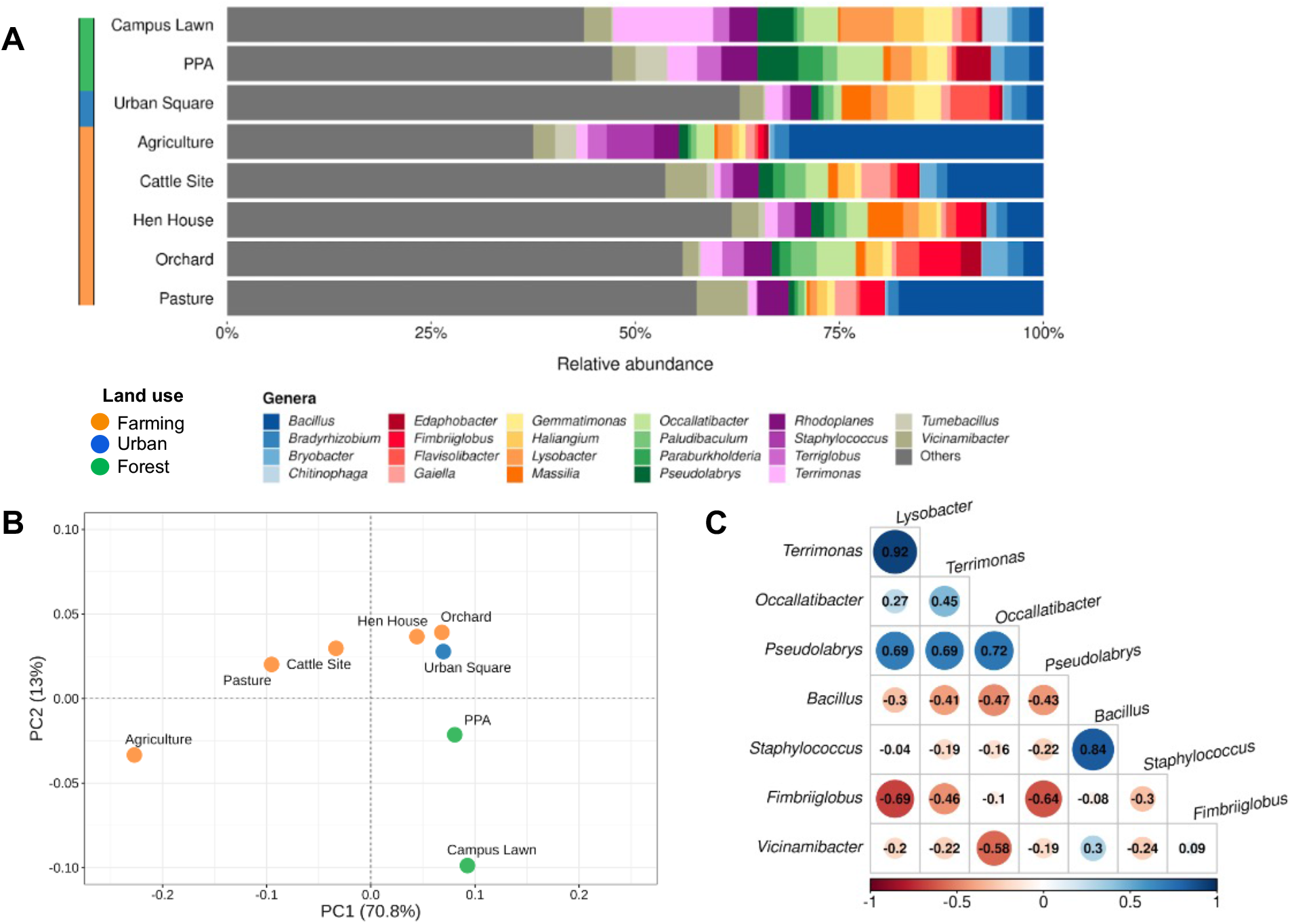
Most abundant taxa and their distribution throughout the studies sites. **(A)** Relative abundances of groups found in environmental samples according to 16S sequencing. All taxa found with relative abundances below 2.5% in each sample were labeled as “Others”. **(B)** Principal component analysis (PCA) of most abundant bacteria, clustered together per sampling site and colored according to land use classification. **(C)** Correlogram indicating negative (in red) and positive (in blue) correlations among most abundant taxa (minimum relative abundance of 5%) in the studied sites.

Additionally, in order to comprehend bacterial diversity in each area, the Shannon index was calculated for each soil sample, indicating a smaller diversity in agricultural soils (3.781), followed by campus lawn (4.343) and PPA (4.504), and the highest diversity found in urban square (5.056) and hen house soils (5.087), in the analyzed conditions (Table S3). We next performed a principal component analysis (PCA) and a hierarchical clusterization (Figure 1B and Figure S2, respectively) with the resulting microbial profiles. The results of both analyses corroborated with the land use classification used here.

To further understand the dynamics of bacterial communities, we performed a correlation analysis for taxa with a minimum relative abundance of 5% (Figure 1C). As shown in the figure, *Terrimonas* and *Lysobacter* displayed the strongest positive correlation (0.92), followed by *Bacillus* and *Staphylococcus*. On the other hand, *Fimbriiglobus* and *Lysobacter* displayed the strongest negative correlation (−0.69), followed by *Fimbriiglobus* and *Pseudolabrys* (−0.64) and *Vicinamibacter* with *Occallatibacter* (−0.58).

### 3.2. Abundance and diversity of ARGs and VFs in soils’ metagenomes

Aiming to understand the abundance and diversity of antibiotic resistance genes (ARGs) and virulence factors (VFs) across studied samples, we proceed to analyze the soil metagenomes based on shotgun sequencing through the Illumina platform (Figure 2). Using the CARD database, we detected ARGs in all sampled sites with a total of 254 ARGs identified, ranging from 17 (PPA) to 60 (Cattle site) (Table S5). The identified ARGs can confer potential resistance to eight pharmacological classes of antibiotics, being glycopeptide (77.5%), rifamycin (12.2%) and macrolide/penam (5.9%) the most frequent ARG types across all soils analyzed, followed by trimethoprim (1.1%), phenicol (0.8%), aminoglycoside (0.4%) and isoniazid/rifamycin (0.4%) (Figure 2A).

**Figure 2.**
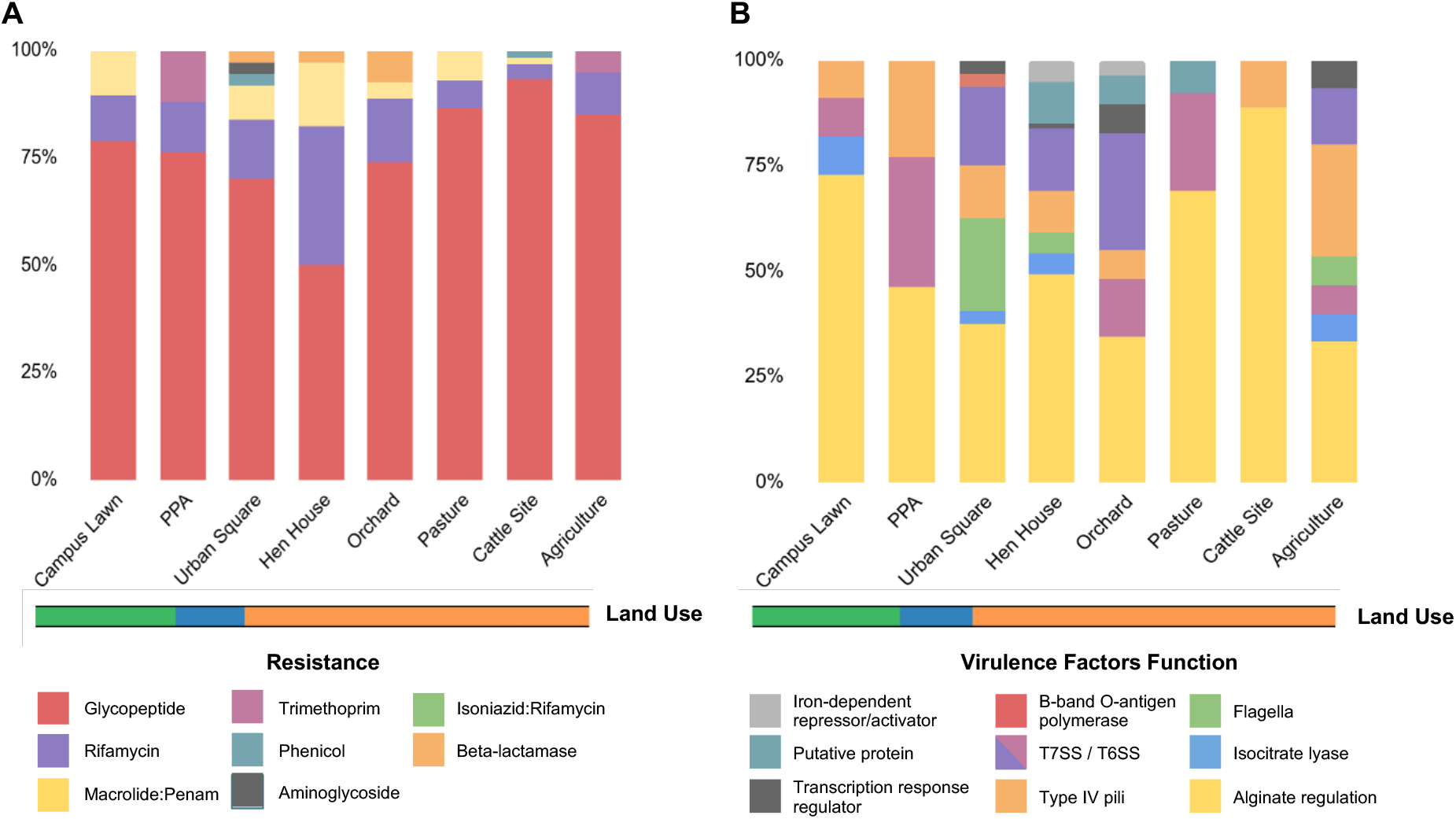
Relative abundances of ARGs and VFs per soil metagenome. (A) Relative abundances of the identified ARGs per soil metagenome, colored by resistance category, as indicated in the legend. (B) Relative abundances of the identified VFs per soil metagenome, colored by function, as indicated in the legend. The ARGs were identified by CARD and VFs by VFDB and the abundances were estimated by dividing the number of distinct resistance genes in the category (i.e., ARG potential resistance or VF function) by the total number of genes for all classes found in that site. Land use is indicated in orange (Farming), blue (Urban) and green (Forest).

In total, 12 ARG types were identified in all soils, being *vanRO* and *vanSO* (glycopeptide)*, rbpA* (rifamycin) and *mtrA* (macrolide/penam) the most frequent genes, followed by *dfrB3* and *dfrB7* variants (trimethoprim), *cpt* (phenicol), *aac2-lb* (aminoglycoside) and *efpA* (isoniazid/rifamycin resistance). Additionally, three β-lactamase genes were identified, being one a serine-β-lactamase (SBL) identified as *blaF* in urban square soils and two metallo-β-lactamase (MBL) encoding genes, identified as *bla*BJP-1 and *bla*LRA-9 in orchard soils (Figure 3A).

**Figure 3.**
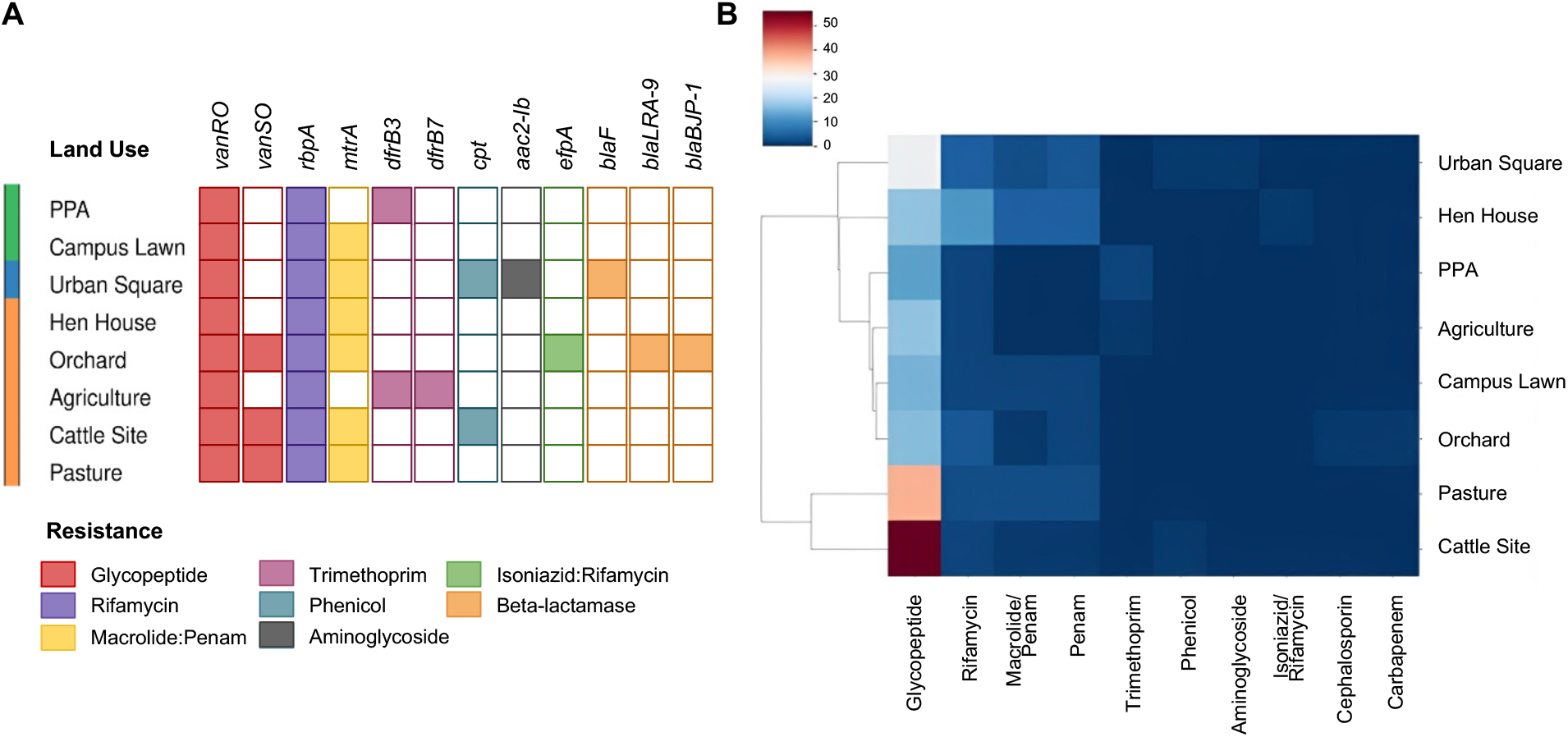
Identification of ARGs in sequenced soil samples. (A) Distribution of different ARGs per soil metagenome, colored by resistance category, as indicated in the legend. (B) Heatmap showing the presence of ARGs identified by CARD indicating the number of each resistance gene per soil. Data were clustered using hierarchical mapping with Euclidean distance. The blue to red scale indicates the number of ARGs for each soil sample in each category. Land use is indicated in orange (Farming), blue (Urban) and green (Forest).

Divergently to what was observed on the dendrogram based on microbial composition, the clusterization of soils based on ARGs’ frequencies indicated two distinct groups, namely, (i) livestock soils (pasture and cattle sites), which are composed of *vanRO* and *vanSO, rbpA* and *mtrA* and (ii) all other soils, in which a higher diversity of ARGs could be identified, especially in urban square and orchard soils, including, in addition to the aforementioned β-lactamases genes, the two *dfrB* variants, *cpt*, *aac2-lb* and *efpA* ARG types (Figure 3B).

To visualize connections between the resistome (dis)-similarities of the selected soils under different land uses, ARGs’ relative abundances across the studies sites were used to plot a chord diagram (Figure 4A), which indicates that even though three ARGs are widespread in all soils, namely *vanRO, mtrA and rbpA*, a higher abundance of genes in cattle sites and pasture soils is observed, whilst the highest diversity of ARGs was identified in orchard and urban square soils.

**Figure 4.**
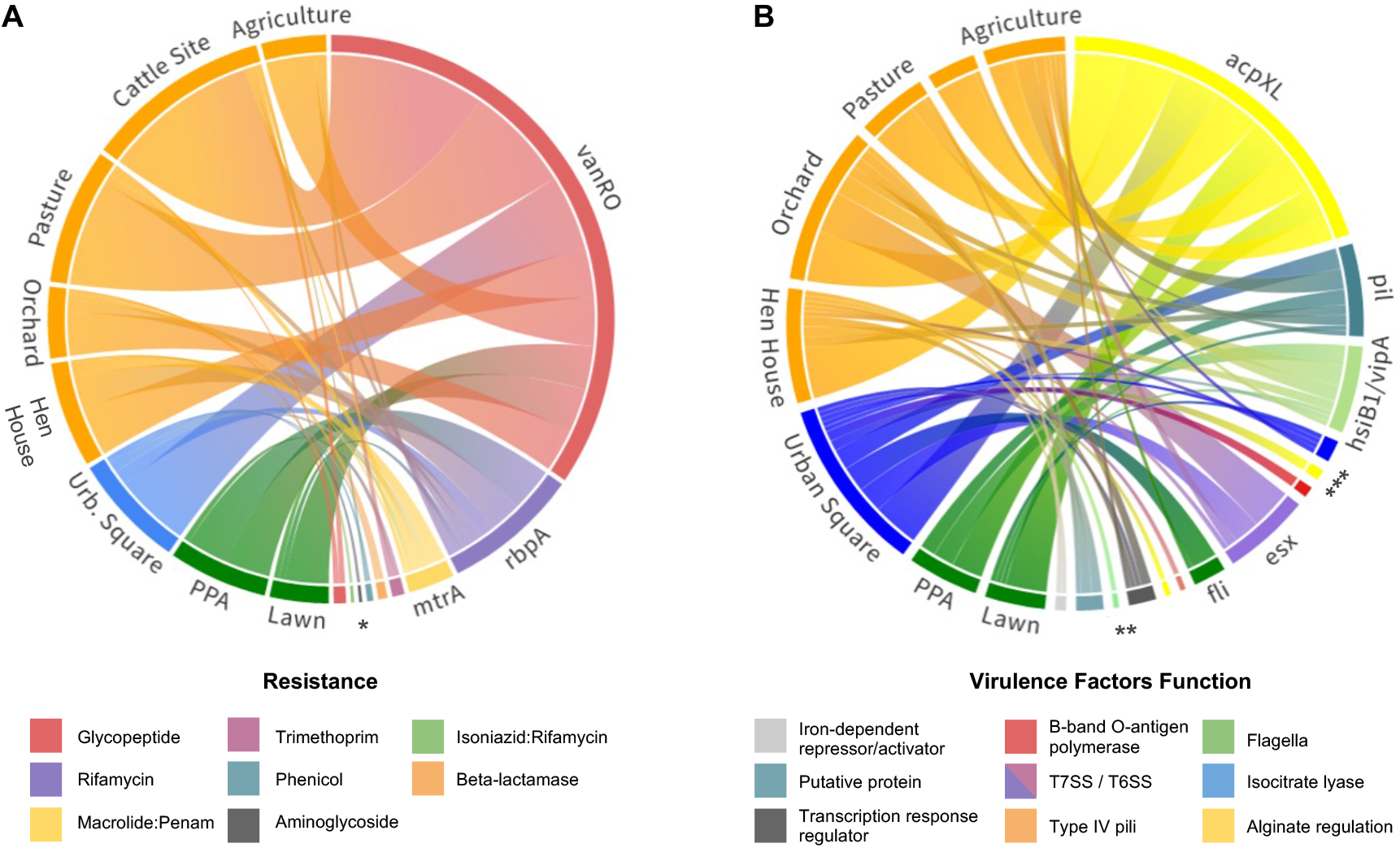
Identified gene distribution across sequenced soil samples. (A) Chord diagram showing the distribution of different ARGs per soil metagenome. Resistance genes are colored by resistance category and soil sites are colored by land use system as follows: orange - farming; blue - urban; green - forest. (B) Chord diagram showing the distribution of different VFs per soil metagenome. Virulence-related genes are colored by function, as indicated in the legend. (*acpXL, alg, mucD*) Alginate regulation; (*icl*) Isocitrate lyase; (*hsiB1/vipA*) Type VI secretion system, T6SS; (*flg, fli, cheW*) Flagella; (*pil*) Type IV pili; (*esx*) Type VII secretion system, T7SS; (*waaG*) B-band O-antigen polymerase; (*phoP*) Possible two component system response transcriptional positive regulator; (*mbtH*) Putative protein; (*ideR*) Iron-dependent repressor and activator. * (*vanSO, efpA, aac2-lb, cpt, bla, dfrB*); **(*ideR, mbtH, phoP, mucD, waaG*); ***(*flgC, alg, icl*). Resource: https://public.flourish.studio/visualisation/116771 & https://public.flourish.studio/visualisation/116970.

From the VFDB database, 143 virulence genes were detected in all soil samples, divided into 25 different genes assigned to 10 classes referring to their virulence functions (Figure 2B). Among them, the main virulence factor was an acyl carrier protein encoded by the *acpXL* gene, that corresponds to 45.5% of all VFs identified throughout the soils and whose main function includes adding very long-chain fatty acid (VLCFA) to lipid A, the hydrophobic group of LPS in Gram-negative bacteria [35]. Nonetheless, types IV and VI secretion systems together compose 22.4% of the virulence genes found (Table S6-S8), mostly involved in adaptation and manipulation of their environment and also in the aggravation of infectious conditions when present in pathogenic bacteria [36]. Nonetheless, genes associated with bacterial motility related to type IV pili, such as *pilT, pilM, pilG* and *pilH*, and rotating flagella, such as *flgC, fliE, fliQ, fliP, fliN* and *fliA* were also widespread in soils. Additionally, when aligning our sequences against the PlasmidFinder database, no plasmid markers were identified in our metagenomic data.

## 4. Discussion

### 4.1. Bacterial Community Structure and Dynamics

The clusterization of soils based on microbial composition on our analyses corroborated with the land use classification previously used, which might indicate that bacterial communities in soils are shaped and modified according to land use over the years, endorsing previous reports in the literature [37–39]. *Vicinamibacter* (Acidobacteriota) and *Rhodoplanes* (Pseudomonadota) genera were ubiquitous and abundant in all soils, likely due to their essential roles in carbon, nitrogen and sulfur cycling [40–42]. Forest soils presented smaller diversity indexes compared to urban and livestock soils (Table S3), which corroborates with previous hypothesis that higher taxonomic diversity is essential to stressed soils maintenance [43]. The main groups found in those soils were Pseudomonadota (*Lysobacter, Pseudolabris* and *Bradyrhizobium*), Acidobacteriota (*Ocallatibacter*) and Bacteroidota (*Terrimonas*) [44–46], with a smaller abundance of *Bacillus* (Bacillota) species compared to other sites.

In the other hand, urban soils presented, along with hen house soils, the highest diversity indexes, which could be explained by the accumulation of human activity wastes and by-products in the former as well as the introduction of gastrointestinal microbiota members in the latter [47]. Both soils were abundant with *Massilia* (Pseudomonadota) members, common environmental bacteria that have been shown to cause opportunistic infections in immunocompromised patients [48,49]. Farming soils presented a higher abundance of *Bacillus*, especially in livestock and agricultural soils, indicating a dominance of this group in farming land use systems in São Paulo’s northeastern soils, corroborating previous reports [50,51]. This could be explained by spore-forming characteristic of *Bacillus*, which facilitates their high resistance to most adverse environmental conditions on farming land use systems, such as heat, desiccation and high levels of UV radiation [44,51].

In agricultural soils, *Staphylococcus* and *Bacillus* represented the majority of sequenced members (Figure 1C), which could be connected to a positive correlation of 0.84 observed between both genera. As expected, agricultural soils presented reduced microbial diversity compared to other soil samples, which could be attributed to soil microbial community homogenization due to the intensified land use of sugarcane crop soils [52,53], such as the one collected for this study.

Some environmental microorganisms, such as *Staphylococcus* and *Bacillus* identified in the studied soils, not only are commonly found in the environment due to their important interactions with other bacteria and functional-maintenance in soils, but also can infect humans and other animals [54]. This ubiquitous characteristic of these microbes is likely due to their high fitness to colonization and proliferation on plant and soil surfaces, related to their high nutritional competition potential and antimicrobial metabolites production, suggesting a possible transfer mechanism to human hosts, especially through the food chain [55]. An imbalance caused by anthropic activities on soil microbial communities could favor *Staphylococcus* and *Bacillus* species [56], in which their biodegradability potential could increase their interactions and stimulate an early proliferation, taking advantage of transient conditions to outgrow more fastidious opportunistic microorganisms [55,57]. It is important to highlight that members of the genus *Bacillus* are one of the most abundant bacterial genera found in soils, with species widespread in a wide range of ecological niches, which is due to their genetic and metabolic diversity, allowing them to serve multiple ecological functions in the environment, including nutrient cycling and conferring abiotic and biotic tolerance to plants [59]. Along with their ecological roles, some members of the genus can be responsible for human infections. *Bacillus anthracis* for instance, can be found in soils as endospores and might be inhaled by humans under certain climatic conditions, resulting in anthrax disease, while *Bacillus cereus* act as a commensal probiotic bacteria resident in gastrointestinal tract, though studies indicated that virulent opportunistic strains can cause not only local infections, such as gastroenteritis, but also hemolytic effects on vertebrates, due to their toxic proteins production [60,61].

*Staphylococcus* spp. are common in the biosphere and possess the capacity to withstand extreme temperatures and pH variations, allowing them to colonize different habitats [62]. Due to its high infectivity capacity, *Staphylococcus aureus* is a well studied commensal and opportunistic species able to colonize a wide variety of hosts and environments, being able to express several toxins associated with distinct disease, such as toxic shock syndrome, along with the biofilm formation capacity, enhancing the persistence of infections and causing resistance towards several drugs [63,64]. This pathogenic bacterium is also responsible for outbreaks of nosocomial and community-associated infections due to methicillin-resistant *S. aureus* (MRSA), which has been reported worldwide [65,66].

### 4.2. Resistance Genes and Virulence Factors Identification

The soil clusterization based on ARGs’ frequencies indicated that, although *vanRO*, *mtrA* and *rbpA* were widespread through all sampled soils, the livestock soil resistomes shared a closer relationship, due to their high abundance and small diversity of ARGs, in comparison to other soils, in which a smaller abundance of genes was detected. Both *vanRO* and *vanSO*, components of the same *vanO* gene cluster that can potentially confer glycopeptide resistance, were firstly identified in *Rhodococcus equi* soil isolates [67]. Glycopeptide antibiotics, such as vancomycin, are a last-resort treatment option for MRSA and enterococci infections [68,69]. In Brazil, several waves of resistance of *S. aureus* against antimicrobials have been reported, being the second and the fourth waves related to hospital- and community-acquired MRSA, respectively, with increasing numbers of resistant strains isolated in different hospitals in São Paulo [70,71]. Recent reports have suggested a concerning scenario, indicating that all Brazilian regions have notified the presence of MRSA strains, especially in the southeastern region, the most affected in the last decades [71]. Although vancomycin resistance genes are commonly found in soils [72,73], our findings highlight the importance of environmental surveillance, since *Staphylococcus* were one of the major components of agricultural soils, abundant with *vanRO* genes.

Macrolide antibiotics act by binding to the bacterial 50S ribosomal subunit causing the cessation of bacterial protein synthesis, as a bacteriostatic agent [74]. The broad antibacterial activity of this antimicrobial has led to their widespread use in gastrointestinal, respiratory tracts and in sexually transmitted infections [74]. In staphylococcal infections, there is an increasing cross-resistance to macrolides in MRSA strains, categorizing these bacteria as a pathogen of great concern, especially due to the high mortality rates and long hospitalization periods related to those [75]. β-lactam antibiotics share the presence of a β-lactam ring in their structures, having activity against a broad spectrum of bacteria due to the penicillin-binding protein (PBP) inactivation, responsible for the cell wall formation [76]. These are the most prescribed antibiotic classes in clinical settings worldwide, with annual expenses of approximately US$ 15 billion, representing 65% of the total antibiotic market [77,78]. In the last few years, the dissemination of Gram-negative bacteria resistant to β-lactams has been considered a public health threat, especially when examining that in the last 20 years, only two new antibiotics have been developed, none with activity against Gram-negative bacteria [79]. The transcriptional activator of the *mtrCDE* multidrug efflux pump, *mtrA* (widespread through all sampled soils of this study), is responsible for, under inducing conditions, expressing the operon that exports a wide variety of antimicrobial agents, including β-lactams and macrolides [80]. The aforementioned gene has been previously reported as abundant in soils, especially in those that have undergone land use conversion [81,82], which could be linked to additional roles for this transcriptional activator in soil bacteria, such as its regulation of antibiotic production in *Streptomyces* [5].

Rifamycin resistance genes have been reported as abundant in both pristine and highly modified soils, which goes in accordance to our findings [83], although few studies have reported such a robust presence of the *rbpA* gene in soils [84]. The genetic product is an RNA-binding protein that has demonstrated to confer low levels of rifamycin resistance in *Streptomyces coelicolor* [85] and it might influence the response of *Mycobacterium tuberculosis* against rifamycin antibiotics [86]. In 2019, 73,000 new cases of tuberculosis (TB) and 4,500 deaths due to this disease have been reported, being several of them related to rifampicin-resistant strains, in a way that TB treatment with 82% chance of success decreases to 60% when associated with isoniazid and rifampicin resistant *M. tuberculosis* strains [87,88]. Additionally, genomic characterization of the zoonotic and human-opportunistic pathogens *Rhodococcus equi* and *Mycolicibacterium peregrinum* obtained from human, pig and soil samples indicated that all the isolates’ genomes contained *rbpA* genes, suggesting that infections caused by these antibiotic resistant bacteria might have an environmental source [89,90]. The presence of *Rhodococcus* bacteria forming a minor component in livestock and hen house microbiotas, *rbpA*-enriched soils, reinforces the need of environmental surveillance.

While fewer genes represent the majority of ARGs in livestock soils, the highest diversity of ARG types was identified in urban square and orchard soils, including β-lactamase-encoding genes, whose products might confer resistance to most of the drugs included in the β-lactam class, which correspond to the vast majority of less toxic options used to treat bacterial infections [76]. These enzymes are capable of inactivating β-lactam antibiotics and can be classified either as SBLs, with an active site containing a catalytic serine residue [91], or as MBLs, that use zinc as cofactor to catalyzes [92].

The two MBL-encoding genes (*bla*LRA-9 and *bla*BJP-1) identified in orchard soils are categorized in the B3 MBL subclass and were previously reported on environmental samples [93,94], conferring high levels of resistance when expressed in *Escherichia coli* clones [95].

Although no reports, to our knowledge, of the aforementioned MBLs have been performed on clinical settings, the occurrence of these genes in orchard soils could pose a threat to human health if they migrate to pathogens. For example, in the case of *bla*BJP-1, which confer less sensibility to chelating agents compared to other MBLs [91] and a high catalytic activity with meropenem – a Watch group antibiotic. Thus, transfer of this ARGs could lead to a risk of selection of bacterial resistance that should be prioritized as targets of stewardship programs and monitoring [96]. Carbapenem antibiotics have a broad activity spectrum against Gram-negative, Gram-positive and anaerobic bacteria, with great performance to extended-spectrum β-lactamases, but may be more susceptible to MBLs [97]. Although intrinsic carbapenem resistance is presented by some bacterial species, by the production of endogenous MBLs, acquired resistance, resulted by horizontal gene transfer, is more common in clinically important bacteria, which highlights the potential thread related to the presence of MBLs in the studied soils [78].

The SBL-coding gene identified, *blaF*, is a chromosomally encoded class A β-lactamase and has been previously correlated with *Mycobacterium fortuitum* (Actinomycetota), a nontuberculous mycobacterium that is commonly found in soils and causing opportunistic infections in humans [98, 99]. It presented a broad-spectrum activity against most β-lactam antibiotics, with the exception of third-generation cephalosporins [100,101]. Few studies have indicated the presence of this β-lactamase in soils, reinforcing the need of surveillance, aiming to report, characterize and highlight further information regarding environmental β-lactamases.

Considering the results related to virulence factors identification in the eight metagenomes, the *acp*XL gene was found in higher prevalence and it encodes an acyl carrier protein, required in the process of adding very long-chain fatty acid (VLCFA) to lipid A, which is a modification performed to preserve the outer membrane barrier function, increasing the hydrophobicity of the bacterial cell LPSs (liposaccharides) and in its rigidity [35,102]. LPSs are known for their role in bacterial invasion, an essential function for the host infection process, and in bacterial adaptation in the environment, regardless of established mutualistic or pathogenic interactions [103]. The VLCFA attached to lipid A has been found in most Rhizobiaceae, as well as in *Bradyrhizobium*, both found in the soils of our study, where the stability function added to the external membrane by its presence could confer greater tolerance to stress and adaptation in several habitats [35].

Bourassa et al. (2017) [102] held studies with mutant *acp*XL strains to determine the structure of LPS lipid A and to determine symbiotic and stress tolerance phenotypes, in which they obtained results that determined that mutants for the gene presented alteration of stability properties of the membrane, altered sensitivity to cationic peptides, in addition to decreased levels of osmoprotective cyclic β-glucans. In addition to this function of defense against stress, VLCFA can also be found in pathogenic or intracellular strains, such as the pathogen *Brucella*, for example, in which this linked lipid A ensures poor recognition of the pathogen’s lipid A by innate immunity, important for the stealth mode mechanism of virulence employed by these bacteria [104].

Protein secretion systems, composing the second most common virulence class found in our study, are used for bacterial cells to interface with their environment through interaction and manipulation, where the secreted proteins can act as virulence factors, allowing these interactions 105]. The type VI secretion system (T6SS), with relative abundances of 11.2% in our samples, is established by the VipA protein. This complex acts as a specialized bacterial nanomachinery that either translocates protein particles to the interior of target cells or releases them to the environment, functioning as a widespread and effective instrument by which bacteria can interact with abiotic environments, eukaryotic and other prokaryotic cells. Thus, it consequently acts as an important determinant of the pathogenicity of eukaryotic cells, as well as in their competitive fitness in the community, since unfit members of microbiota are commonly led to death or growth arrest [106,107]. This indicates that secretion systems have a key role in shaping the microbiota of many ecological niches, changing population ratios of the communities, not only through toxins targeting vital functions, but also leading to starvation due to the prevention of nutrient uptake in competitor bacteria [108,109]. The TSS6, which comprehended 11.2% of all VF hits, could be increased by the acquisition of individual effector modules, as well as small operons, or in the case of Pseudomonadota via horizontally acquired T6SS operons on integrative conjugative elements.

The second VF with the highest relative abundance was secretion system type VII (T7SS). This system is a specialized protein secretion machinery that transports substrates through the cell envelope, widespread in Gram-positive representatives of Actinobacteriota and Bacillota phyla, with effectors capable of affecting a number of bacterial processes, such as sporulation, conjugation, and cell wall stability [107]. Nonetheless, it is believed that the small virulence proteins secreted by T7SS are important for housekeeping functions, such as bacterial cell wall maintenance, seen they’re widespread among pathogenic and environmental microorganisms [107, 110].

Bacteria, in order to interact with external environmental stimuli, exhibit long proteinaceous appendages on their cell surface in a wide range of bacteria, determined as pili or fimbriae, as identified in our targeted soils [111,112]. These non-flagellar structures are classified according to their assembly pathways, being type VI pile, found predominantly in our study, whose flexible surface appendages act as virulence factors in Gram-negative bacteria, and type IV-related pili, that are also identified in Gram-positive, capable of driving environmental adaptation and fitness [113,114].

It’s important to note that there is great diversity among the structures and functions of these flagella, which serve not only as motility machines but also biosensors to solid surfaces, facilitating the lifecycle transition, along with biofilm formation mediation, which in the environment has been associated with greater protection against desiccation and toxic compounds, in addition to the retention of matrix enzymes that improve the efficiency and diversity of the decomposition of the organic matter [115–117].

## 5. Conclusions

The concept of One Health highlights that human health is interconnected to the health of other members on ecosystems, such as soils, animals and plants. In that sense, microorganisms are crucial in one health, since they are the links among all these members, seen in the role of commensal bacteria on driving the organisms’ fitness, as well as maintaining key soil functions. Here, we showed that structure and composition of the microbial communities of soil samples correlates to its land use, along with concerning interactions among environmental and opportunistic pathogens in soils that have undergone land conversion. Our study has also shown several commonly found ARGs in soils that are also responsible for antibiotic resistance in potential pathogenic bacteria widespread in soils, with the highest abundances present in livestock soils, providing the first, to our knowledge, environmental AMR surveillance of soil samples in a crucial region of Brazil in terms of population density and economic relevance. Although ARGs found in the soil samples in this study may confer resistance against competitors in these habitats, their gene products may also serve other functions in soils. We reinforce that ARG or VF hits within samples don’t mean actual antibiotic resistance or actual virulence determinants. Nevertheless, identifying MBLs in highly modified soils highlights the importance of environmental surveillance to pull the brakes and gather more information regarding resistance levels in regions at risk for higher selective pressure due to anthropic activities.

## Authors Contributions

Contributed to the conception and design of the study – JVWO, AFTF and MEG. Sample collection and processing – JVWO and AFTF. Performed the wet-lab procedures – JVWO and SV. Bioinformatic data analysis (amplicon and shotgun, respectively) – GT and RSR. Manuscript writing and paper design - JVWO, GMS and MEG. Review and approval of the final version of the paper – all authors. All authors contributed to the article and approved the submitted version.

## Data Availability Statement

Soil metagenomes were deposited in the SRA repository under the BioProject number PRJNA900430 (Table S4).

## Acknowledgments

The authors thank their laboratory colleagues for their insightful comments and suggestions throughout this study. The authors also would like to thank the lab technician Thalita Riul Prado for her assistance in the course of this work and to Guilherme Marcelino Viana de Siqueira for his initial support on microbiota data analysis.

