## Supplemental Figure S1 and S2; Tables S1 to S8 for "Metagenomic insights for antimicrobial resistance surveillance in soils with different land uses in Brazil"

### Supporting Materials

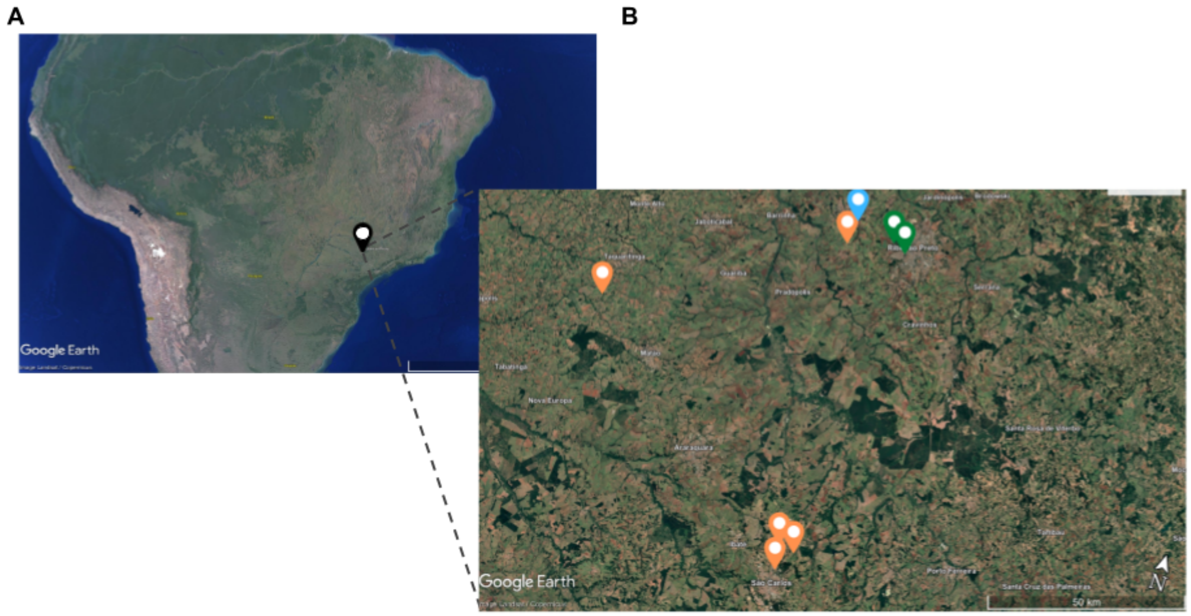


**Figure S1.** Map of soil sample locations. **(A)** Brazil map, with Ribeirão Preto marked in green. **(B)** Part of the São Paulo northeastern region, in which collection sites, along with their land use classification, are shown with green (forest), blue (urban) and orange (farming soils) markers. The map was taken using Google Maps and Google Earth Pro (https://www.google.cl/maps; accessed on 29 July 2022).

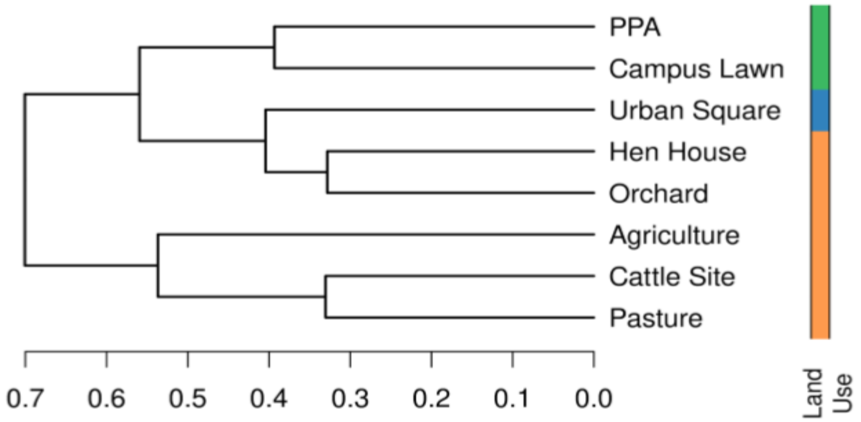


**Figure S2.** Soil sample clusterization based on microbial composition. The cladogram was constructed using Ward's algorithm for hierarchical clustering on a distance matrix obtained with Bray-Curtis dissimilarity method. Land use is indicated in orange (Farming), blue (Urban) and green (Forest).

**Table S1.** Shotgun sequencing - Illumina NovaSeq 6000- data quality summary.

| Sample ID* | Raw Reads | Raw data (G) | Effective (%) | Error (%) | Q20 (%) | Q30 (%) | GC (%) |
|---|---|---|---|---|---|---|---|
| SOrch | 117562592 | 17.6 | 99.87 | 0.03 | 97.36 | 92.70 | 62.79 |
| SLawn | 95499316 | 14.3 | 99.84 | 0.03 | 97.73 | 93.48 | 61.67 |
| SPas | 107762684 | 16.2 | 99.86 | 0.03 | 97.53 | 93.22 | 66.09 |
| SUrbSq | 120088348 | 18.0 | 99.84 | 0.03 | 97.57 | 93.23 | 65.32 |
| SCatt | 124484298 | 18.7 | 99.84 | 0.03 | 97.58 | 93.22 | 66.34 |
| SHen | 104840520 | 15.7 | 99.82 | 0.03 | 97.44 | 92.89 | 65.26 |
| SPpa | 83080300 | 12.5 | 99.82 | 0.02 | 98.04 | 94.51 | 63.29 |
| SAgri | 81025724 | 12.2 | 99.80 | 0.03 | 97.90 | 94.13 | 64.14 |

* SOrch: Orchard; SLawn: Campus Lawn; SPas: Pasture; SUrbSq: Urban Square; SCatt: Cattle Site; SHen: Hen House; SPpa: Permanent Protected Area; SAgri: Agriculture.

**Table S2.** Relative abundances of most abundant genera (above 2.5%) identified in sequenced soils.

| Genus | Soil Sample | Read Count | Relative Abundance |
|---|---|---|---|
| *Occallatibacter* | PPA | 1667 | 0.056 |
| *Pseudolabrys* | PPA | 1483 | 0.050 |
| *Rhodoplanes* | PPA | 1302 | 0.044 |
| *Edaphobacter* | PPA | 1224 | 0.041 |
| *Tumebacillus* | PPA | 1152 | 0.039 |
| *Terrimonas* | PPA | 1083 | 0.037 |
| *Paraburkholderia* | PPA | 890 | 0.030 |
| *Bradyrhizobium* | PPA | 888 | 0.030 |
| *Terriglobus* | PPA | 881 | 0.030 |
| *Vicinamibacter* | PPA | 848 | 0.029 |
| *Lysobacter* | PPA | 764 | 0.026 |
| *Terrimonas* | Campus Lawn | 758 | 0.122 |
| *Lysobacter* | Campus Lawn | 406 | 0.066 |
| *Pseudolabrys* | Campus Lawn | 274 | 0.044 |
| *Occallatibacter* | Campus Lawn | 259 | 0.042 |
| *Haliangium* | Campus Lawn | 229 | 0.037 |
| *Gemmatimonas* | Campus Lawn | 216 | 0.035 |
| *Rhodoplanes* | Campus Lawn | 212 | 0.034 |
| *Vicinamibacter* | Campus Lawn | 207 | 0.033 |
| *Chitinophaga* | Campus Lawn | 192 | 0.031 |
| *Flavisolibacter* | Urban Square | 1684 | 0.048 |
| *Massilia* | Urban Square | 1260 | 0.036 |
| *Gemmatimonas* | Urban Square | 1157 | 0.033 |
| *Haliangium* | Urban Square | 1154 | 0.033 |
| *Vicinamibacter* | Urban Square | 1021 | 0.029 |
| *Rhodoplanes* | Urban Square | 914 | 0.026 |
| *Bacillus* | Agriculture | 335 | 0.312 |
| *Staphylococcus* | Agriculture | 62 | 0.058 |
| *Rhodoplanes* | Agriculture | 33 | 0.031 |

| | | | |
|---|---|---|---|
| *Vicinamibacter* | Agriculture | 29 | 0.027 |
| *Tumebacillus* | Agriculture | 28 | 0.026 |
| *Bacillus* | Hen House | 1833 | 0.044 |
| *Massilia* | Hen House | 1808 | 0.044 |
| *Vicinamibacter* | Hen House | 1352 | 0.033 |
| *Fimbriiglobus* | Hen House | 1260 | 0.030 |
| *Occallatibacter* | Hen House | 1074 | 0.026 |
| *Bacillus* | Pasture | 7529 | 0.177 |
| *Vicinamibacter* | Pasture | 2643 | 0.062 |
| *Rhodoplanes* | Pasture | 1618 | 0.038 |
| *Fimbriiglobus* | Pasture | 1288 | 0.030 |
| *Gaiella* | Pasture | 1119 | 0.026 |
| *Bacillus* | Cattle Site | 3017 | 0.118 |
| *Vicinamibacter* | Cattle Site | 1304 | 0.051 |
| *Gaiella* | Cattle Site | 907 | 0.035 |
| *Rhodoplanes* | Cattle Site | 805 | 0.031 |
| *Occallatibacter* | Cattle Site | 701 | 0.027 |
| *Paludibaculum* | Cattle Site | 653 | 0.025 |
| *Fimbriiglobus* | Cattle Site | 648 | 0.025 |
| *Fimbriiglobus* | Orchard | 996 | 0.051 |
| *Occallatibacter* | Orchard | 949 | 0.048 |
| *Rhodoplanes* | Orchard | 666 | 0.034 |
| *Bryobacter* | Orchard | 614 | 0.031 |
| *Paludibaculum* | Orchard | 612 | 0.031 |
| *Flavisolibacter* | Orchard | 553 | 0.028 |
| *Terrimonas* | Orchard | 522 | 0.027 |
| *Terriglobus* | Orchard | 520 | 0.026 |
| *Edaphobacter* | Orchard | 503 | 0.026 |

**Table S3.** Alpha diversity indexes of sampled soils.

| Soil | Shannon Index | Simpson Index |
|---|---|---|
| Hen House | 5.087 | 0.9867 |
| Urban Square | 5.056 | 0.9866 |
| Orchard | 4.859 | 0.9835 |
| Cattle Site | 4.791 | 0.9746 |
| Pasture | 4.543 | 0.9570 |
| PPA | 4.504 | 0.9783 |
| Campus Lawn | 4.343 | 0.9670 |
| Agriculture | 3.781 | 0.8932 |

**Table S4.** Data availability. The trimmed sequences, along with their quality scores, resulting from shotgun sequencing are publicly available under the BioProject PRJNA900430.

| BioSample Accession | Sample Name/SPUID | Organism | Tax ID | BioProject | SRA Accession |
|---|---|---|---|---|---|
| SAMN31691838 | SHen | soil metagenome | 410658 | PRJNA900430 | SRR22278232 |
| SAMN31691839 | SCatt | soil metagenome | 410658 | PRJNA900430 | SRR22278231 |
| SAMN31691840 | SLawn | soil metagenome | 410658 | PRJNA900430 | SRR22278230 |
| SAMN31691841 | SUrbSq | soil metagenome | 410658 | PRJNA900430 | SRR22278229 |
| SAMN31691842 | SPast | soil metagenome | 410658 | PRJNA900430 | SRR22278228 |
| SAMN31691843 | SOrch | soil metagenome | 410658 | PRJNA900430 | SRR22278227 |
| SAMN31691844 | SAgri | soil metagenome | 410658 | PRJNA900430 | SRR22278226 |
| SAMN31691845 | SPpa | soil metagenome | 410658 | PRJNA900430 | SRR22278225 |

**Table S5.** Summary of the identified ARGs by CARD in sequenced metagenomes of soils under different land use systems.

| ARG | Count | Resistance | Soil | Land Use |
|---|---|---|---|---|
| *vanRO* | 15 | Glycopeptide | Campus Lawn | Forest |
| *rbpA* | 2 | Rifamycin | Campus Lawn | Forest |
| *mtrA* | 2 | Macrolide;Penam | Campus Lawn | Forest |
| *vanRO* | 13 | Glycopeptide | PPA | Forest |
| *dfrB7_1* | 1 | Trimethoprim | PPA | Forest |
| *dfrB3* | 1 | Trimethoprim | PPA | Forest |
| *rbpA* | 2 | Rifamycin | PPA | Forest |
| **Total Forest** | 36 | | | |
| *cpt* | 1 | Phenicol | Urban Square | Urban |
| *aac2-Ib* | 1 | Aminoglycoside | Urban Square | Urban |
| *bla*F | 1 | Beta-lactamase | Urban Square | Urban |
| *vanRO* | 26 | Glycopeptide | Urban Square | Urban |
| *rbpA* | 5 | Rifamycin | Urban Square | Urban |
| *mtrA* | 3 | Macrolide;Penam | Urban Square | Urban |
| **Total Urban** | 37 | | | |
| *mtrA* | 5 | Macrolide;Penam | Hen House | Farming |
| *rbpA* | 11 | Rifamycin | Hen House | Farming |
| *vanRO* | 17 | Glycopeptide | Hen House | Farming |
| *efpA* | 1 | Isoniazid;Rifamycin | Hen House | Farming |
| Total | 34 | | | |
| *blaLRA-9* | 1 | Beta-lactamase | Orchard | Farming |
| *blaBJP-1* | 1 | Beta-lactamase | Orchard | Farming |
| *vanRO* | 15 | Glycopeptide | Orchard | Farming |
| *rbpA* | 4 | Rifamycin | Orchard | Farming |
| *mtrA* | 1 | Macrolide;Penam | Orchard | Farming |

| | | | | |
|---|---|---|---|---|
| *vanSO* | 1 | Glycopeptide | Orchard | Farming |
| Total | 23 | | | |
| *vanRO* | 37 | Glycopeptide | Pasture | Farming |
| *rbpA* | 3 | Rifamycin | Pasture | Farming |
| *vanSO* | 1 | Glycopeptide | Pasture | Farming |
| *mtrA* | 3 | Macrolide;Penam | Pasture | Farming |
| Total | 44 | | | |
| *cpt* | 1 | Phenicol | Cattle Site | Farming |
| *vanRO* | 54 | Glycopeptide | Cattle Site | Farming |
| *rbpA* | 2 | Rifamycin | Cattle Site | Farming |
| *vanSO* | 2 | Glycopeptide | Cattle Site | Farming |
| *mtrA* | 1 | Macrolide;Penam | Cattle Site | Farming |
| Total | 60 | | | |
| *dfrB3* | 1 | Trimethoprim | Agriculture | Farming |
| *rbpA* | 2 | Rifamycin | Agriculture | Farming |
| *vanRO* | 17 | Glycopeptide | Agriculture | Farming |
| Total | 20 | | | |
| **Total Farming** | 181 | | | |

**Table S6.** Summary of the identified VFs by VFDB in sequenced metagenomes of soils under different land use systems.

| VF | Count | Soil | Land Use |
|---|---|---|---|
| *acpXL* | 8 | Campus Lawn | Forest |
| *pilG* | 1 | Campus Lawn | Forest |
| *hsiB1/vipA* | 1 | Campus Lawn | Forest |
| *icl* | 1 | Campus Lawn | Forest |
| **Total** | **11** | | |
| *acpXL* | 6 | PPA | Forest |
| *hsiB1/vipA* | 4 | PPA | Forest |
| *pilG* | 3 | PPA | Forest |
| **Total** | **13** | | |

**Table S7.** Summary of the identified VFs by VFDB in sequenced soil metagenomes on urban land use system.

| VF | Count | Soil | Land Use |
|---|---|---|---|
| *algW* | 1 | Urban Square | Urban |
| *flgC* | 2 | Urban Square | Urban |
| *esxH* | 1 | Urban Square | Urban |
| *fliE* | 1 | Urban Square | Urban |
| *acpXL* | 9 | Urban Square | Urban |
| *hsiB1/vipA* | 1 | Urban Square | Urban |
| *esxN* | 2 | Urban Square | Urban |
| *waaG* | 1 | Urban Square | Urban |
| *mucD* | 1 | Urban Square | Urban |
| *pilT* | 1 | Urban Square | Urban |
| *pilM* | 1 | Urban Square | Urban |
| *fliQ* | 1 | Urban Square | Urban |
| *fliP* | 1 | Urban Square | Urban |
| *fliN* | 1 | Urban Square | Urban |
| *esxM* | 2 | Urban Square | Urban |
| *phoP* | 1 | Urban Square | Urban |
| *icl* | 1 | Urban Square | Urban |
| *pilG* | 1 | Urban Square | Urban |
| *pilH* | 1 | Urban Square | Urban |
| *algR* | 1 | Urban Square | Urban |
| *fliA* | 1 | Urban Square | Urban |
| **Total** | **32** | | |

**Table S8.** Summary of the identified VFs by VFDB in sequenced soil metagenomes on farming land use system.

| VF | Count | Soil | Land Use |
|---|---|---|---|
| *phoP* | 1 | Hen House | Farming |
| *acpXL* | 10 | Hen House | Farming |
| *cheW* | 1 | Hen House | Farming |
| *esxH* | 1 | Hen House | Farming |
| *hsiB1/vipA* | 2 | Hen House | Farming |
| *mbtH* | 2 | Hen House | Farming |
| *ideR* | 1 | Hen House | Farming |
| *pilG* | 2 | Hen House | Farming |
| *icl* | 1 | Hen House | Farming |
| **Total** | **21** | | |
| *phoP* | 2 | Orchard | Farming |
| *mbtH* | 2 | Orchard | Farming |
| *pilT* | 1 | Orchard | Farming |
| *hsiB1/vipA* | 4 | Orchard | Farming |
| *esxM* | 4 | Orchard | Farming |
| *acpXL* | 10 | Orchard | Farming |
| *esxN* | 4 | Orchard | Farming |
| *ideR* | 1 | Orchard | Farming |
| *pilG* | 1 | Orchard | Farming |
| **TOTAL** | **29** | | |
| *acpXL* | 9 | Pasture | Farming |
| *mbtH* | 1 | Pasture | Farming |
| *hsiB1/vipA* | 3 | Pasture | Farming |
| **TOTAL** | **13** | | |
| *pilG* | 1 | Cattle Site | Farming |
| *acpXL* | 8 | Cattle Site | Farming |
| **TOTAL** | **9** | | |
| *acpXL* | 5 | Agriculture | Farming |
| *pilG* | 2 | Agriculture | Farming |
| *esxM* | 1 | Agriculture | Farming |
| *icl* | 1 | Agriculture | Farming |
| *pilT* | 2 | Agriculture | Farming |
| *fliA* | 1 | Agriculture | Farming |
| *esxG* | 1 | Agriculture | Farming |
| *hsiB1/vipA* | 1 | Agriculture | Farming |
| *phoP* | 1 | Agriculture | Farming |
| **Total** | **15** | | |
| **Total Farming** | **143** | | |
